# Gut bacteria mediate nutrient availability *in Drosophila* diets

**DOI:** 10.1101/2020.05.28.120832

**Authors:** Danielle N.A. Lesperance, Nichole A. Broderick

## Abstract

*Drosophila melanogaster* gut microbes play important roles in host nutritional physiology. However, these associations are often indirect and studies typically are in the context of specialized nutritional conditions, making it difficult to discern how microbiome-mediated impacts translate to physiologically relevant conditions, in the laboratory or nature. Here, we show that on three artificial diets and a natural diet of grapes, *D. melanogaster* gut bacteria alter protein, carbohydrates, and moisture of the food substrate. In depth analysis on one diet revealed bacteria also increase tryptophan levels. We investigate how nutrient changes impact life history and find that, while alterations to dietary protein and carbohydrates are arguably the most significant consequence of bacterial association, other factors, such as micronutrients, likely contribute to life history traits in a diet-dependent manner. Our work demonstrates that while some bacterial impacts on nutrition occur across experimental diets, others are dictated by unique dietary environments.

## Introduction

Gut microbes have important functions in host development, immunity, intestinal homeostasis, and metabolism across model organisms (Belkaid and Harrison, 2017; Gerbaba et al., 2017; Lesperance and Broderick, 2020a; Lin and Zhang, 2017). There is growing appreciation for the gut microbiome’s role in host nutritional intake, with effects through both microbial catabolism of nutrients and biosynthesis of metabolites (Chassard and Lacroix, 2013; Ng et al., 2013; Yadav et al., 2018). Use of the *Drosophila melanogaster* model has revealed significant roles for gut microbes in activating nutritional signaling pathways (Shin et al., 2011; Storelli et al., 2011), stimulating protein nutrition (Keebaugh et al., 2018; Yamada et al., 2015), and catabolizing dietary carbohydrates (Huang and Douglas, 2015). A small number of bacterial-derived metabolites, including acetate and B-vitamins, have been identified that promote *D. melanogaster* nutrition either directly or via impacts on feeding behavior (Fischer et al., 2017; Wong et al., 2014). Interactions between host and microbiome are strongly influenced by fly diet. In particular, in conditions of nutrient scarcity, gut bacteria decrease development time and increase lifespan (Keebaugh et al., 2018; Shin et al., 2011; Storelli et al., 2011; Yamada et al., 2015). On the other hand, excess dietary protein is thought to diminish the impact of the microbiome (specifically bacteria) on development and lifespan (Keebaugh et al., 2019).

Despite the many benefits of using *D. melanogaster* to investigate a spectrum of biological processes, the range of diets used for fly maintenance across the field can complicate interpretation (Lesperance and Broderick, 2020b; Lüersen et al., 2019). Recently, we showed that diet composition between studies varied greatly in both the amounts and types of components used (i.e. inactive yeast versus brewer’s yeast versus yeast extract), which ultimately resulted in diets containing a wider than appreciated range of macro- and micronutrients (Lesperance and Broderick, 2020b). Many diets are reported as “standard” in the literature as well, often without basis (for example, protein of so-called “standard” diets analyzed in our study ranged from 6.33 to 77.93 grams per liter), making interpretation of differences in nutritional environments, and, as a result, nutrition-mediated phenotypes difficult (Lesperance and Broderick, 2020b). These discrepancies make it challenging to contextualize studies within and across fields of interest, particularly in the case of nutrition and the microbiome, which depend so heavily on dietary composition.

In this study, we investigate microbial impacts on *D. melanogaster* nutrition by analyzing nutritional content of three “standard” fly diets with and without bacterial inoculation. Two of the diets we tested, the Bloomington Standard and Bloomington Cornmeal-Molasses-Yeast (CMY) diets, are broadly used or are at least the basis of diets used by the *D. melanogaster* community in a range of fly research areas. Thus, they provide a general understanding of microbe-diet relationships on diets in use by a broad swath of the community in contrast to more specialized or defined diets, whose physiologic implications may be difficult to interpret. We assess microbial impacts on fly life history and identify bacteria-mediated changes to protein, carbohydrate, and moisture content, as well as the amino acid tryptophan, as potential mechanisms of microbial modulation of host physiology. We go on to assess nutritional changes due to bacteria on a natural fly diet of grapes to contextualize the overall relevance of our findings.

## Results

### Drosophila gut bacteria impact nutrient content of fly food

To address how the microbiome interacts with diet in laboratory conditions, we prepared three “standard” diets: the Bloomington Standard, Bloomington Cornmeal-Molasses-Yeast (CMY), and our personal laboratory standard diet, called here Broderick Standard. We inoculated bottles of sterile food with either PBS or a bacterial cocktail of four common *D. melanogaster* gut microbes, *Lactobacillus plantarum, Lactobacillus brevis, Acetobacter pasteurianus,* and *Acetobacter tropicalis*, in equal proportions. Treated food was incubated at 25°C for 14 days to simulate the length of time bacteria would associate with food over the course of a typical fly life cycle (from egg laying to adult eclosion). After incubation, food was collected and analyzed for bacterial load and nutritional content including protein, carbohydrates, ash, fat, and moisture. Across the three diets, bacterial treatment decreased carbohydrates and increased moisture compared to PBS-treated controls, with generally no significant change in protein, ash, or fat levels (**Figure 1A-C**, **Figure S1A**). To determine if this relationship between microbes and fly food persisted under a natural method of inoculation, we generated gnotobiotic flies by feeding axenic adults the same 4-species bacterial cocktail, then placed either axenic flies (Sterile treatment) or the gnotobiotic flies (+Bacteria treatment) on sterile food. After 4 days, all flies were removed and food was further incubated for 10 days to allow larvae to pupate and reduce confounding effects of host biomass on nutritional analysis. As food was collected for sampling, emerged adults, larvae, and pupae were discarded. The nutritional values were generally consistent with the levels observed in food directly inoculated with bacterial cultures, though the degree of carbohydrate reduction and moisture increase was greater from gnotobiotic inoculation compared to culture inoculation (**Figure 1E-G, Figure S1B**) despite the genus-level bacterial load being similar for each inoculation method and across diets (**Figure 1D, H**). While protein levels appeared to decrease in the CMY +Bacteria treatments, we noted an associated increase in protein in the Sterile CMY treatment under gnotobiotic conditions as compared to culture-inoculated. Thus, we attribute the difference in protein between Sterile and +Bacteria CMY to possible “contamination” by host tissues, which artificially inflated the Sterile treatment protein levels, since the protein level in the +Bacteria treatments was not different between gnotobiotic and culture inoculation methods.

**Figure 1.**
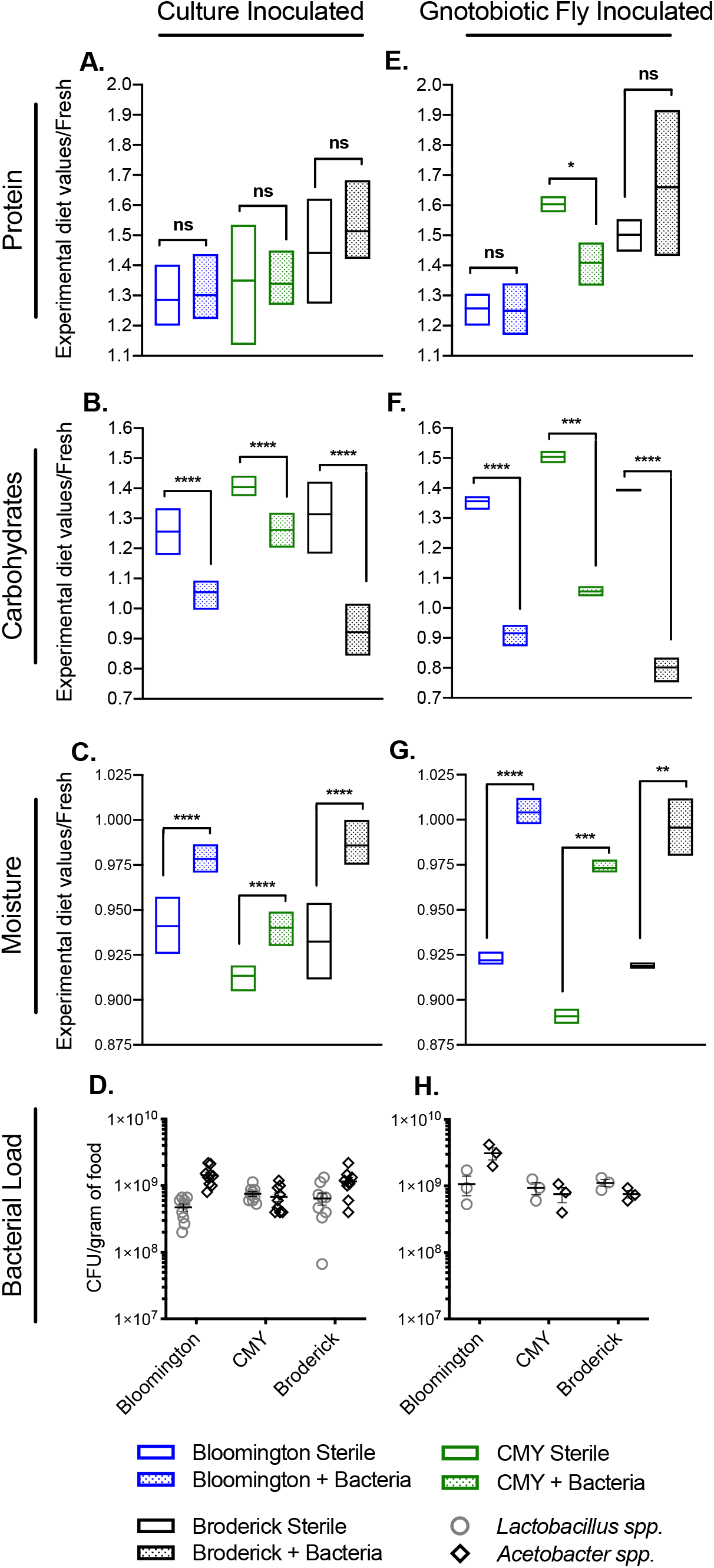
*Drosophila* gut bacteria impact nutrient content of fly food. **A)** Protein, **B)** carbohydrates, **C)** moisture, and **D)** bacterial load in Bloomington, CMY, and Broderick diets 14 days after direct inoculation with PBS (Sterile) or 10^4^ cells of a 4-species bacterial cocktail (+Bacteria). **E)** Protein, **F)** carbohydrates, **G)** moisture, and **H)** bacterial load in Bloomington, CMY, and Broderick diets 14 days after inoculation via axenic flies (Sterile) or gnotobiotic flies previously fed 10^4^ cells of a 4-species bacterial cocktail (+Bacteria, see **Figure S1** for bacterial counts within gnotobiotic flies at time of diet inoculation). Nutritional data **(A-C, E-G)** is expressed as raw values of each nutritional test in treated samples divided by raw values of fresh, untreated diets (see **Figure S1** for raw nutritional data); bars represent minimum and maximum values and mean of 9 **(A-D)** or 3 **(E-H)** biological replicates; statistical differences between Sterile and +Bacteria treatments within each diet were determined using unpaired two-tailed t-tests. Bacterial load **(D, H)** is expressed as colony forming units (CFU) per gram of food with each point representing an individual replicate; lines and error bars represent mean ± SEM; no statistical differences between *Lactobacillus spp.* and *Acetobacter spp.* within each diet and between diets were detected via analysis with two-way ANOVA; bacterial growth was not detected in sterile treatments and is not shown. Significance is expressed as: P>0.05=ns, P≤0.05=*, P≤0.01=**, P≤0.001=***, P≤0.0001=****.

To account for possible differences in food layer depth, we next separately analyzed the nutritional content of the top versus bottom layers of food after treatment with bacteria/PBS and bleached fly embryos. Carbohydrates were decreased and moisture increased in both the top and bottom layers of food, consistent with the whole sample analysis. (**Figure 2A-C**). However, in separating the top and bottom halves of the diet, we observed an increase in protein in the top, but not bottom, layer of food (**Figure 2A**), which was correlated with 10-fold higher bacterial density in the top versus bottom half of food, though bacterial counts of both *Lactobacillus spp.* and *Acetobacter spp.* were significant in both layers (**Figure 2D**). Altogether, this indicates that bacteria are capable of changing the nutritional environment of the food, resulting in relatively low carbohydrates and high moisture throughout, while generating a stratification of protein that correlates with bacterial density.

**Figure 2.**
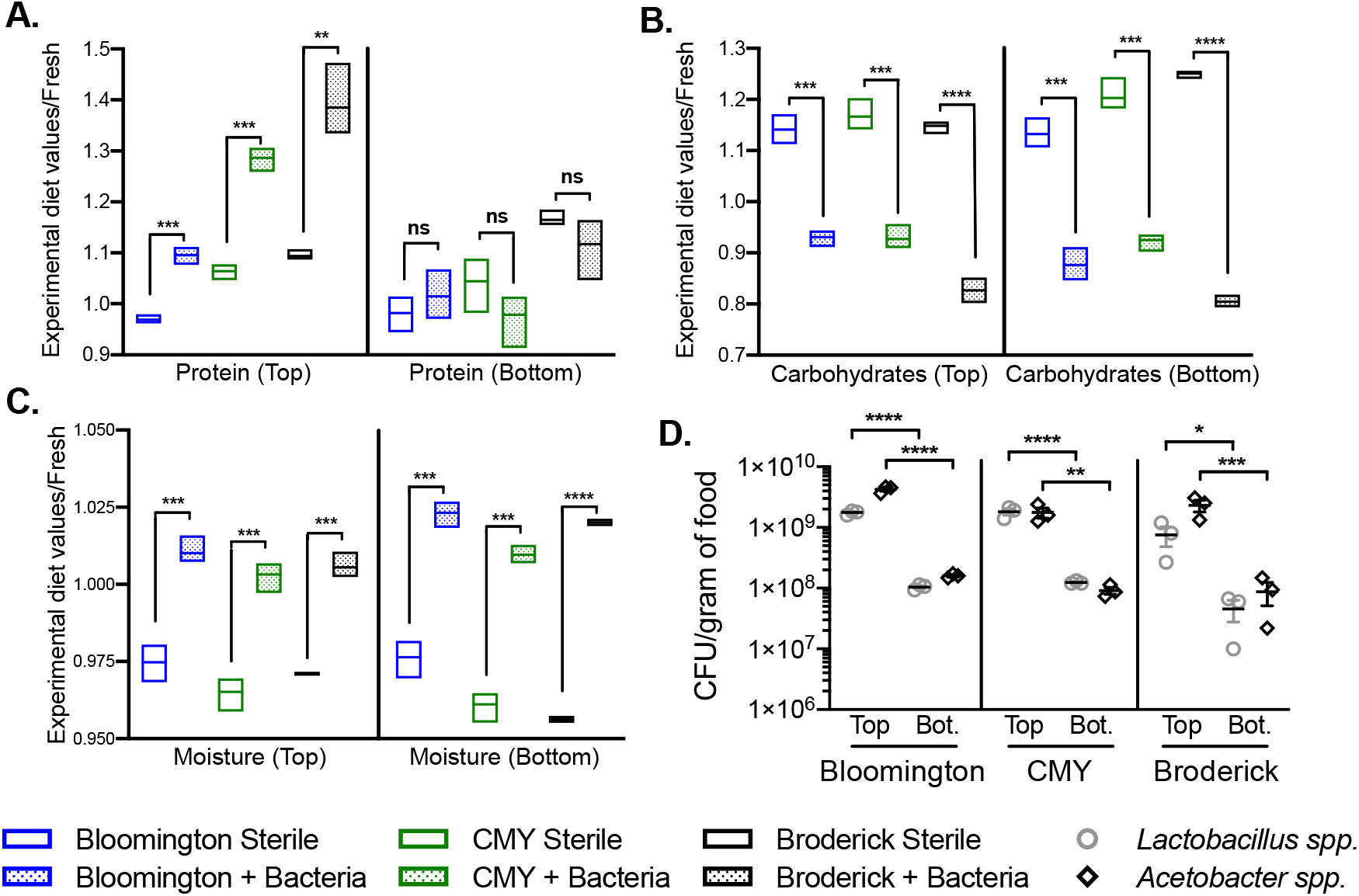
Bacterial growth on fly food results in nutrient stratification. **A)** Protein, **B)** carbohydrates, **C)** moisture, and **D)** bacterial load in Bloomington, CMY, and Broderick diets 11 days after inoculation with sterile embryos and PBS (Sterile) or 10^4^ cells of a 4-species bacterial cocktail (+Bacteria); top and bottom half of food were analyzed as separate samples. Nutritional data **(A-C)** is expressed as raw values of each nutritional test in treated samples divided by raw values of fresh, untreated diets (see **Figure S1** for raw nutritional data); bars represent minimum and maximum values and mean of 3 biological replicates; statistical differences between Sterile and +Bacteria treatments within top or bottom of each diet were determined using unpaired two-tailed t-tests. Bacterial load **(D)** is expressed as colony forming units (CFU) per gram of food with each point representing an individual replicate; lines and error bars represent mean ± SEM; statistical differences between *Lactobacillus spp.* and *Acetobacter spp.* in top versus bottom were detected via two-way ANOVA; bacterial growth was not detected in sterile treatments and is not shown. Significance is expressed as: P>0.05=ns, P≤0.05=*, P≤0.01=**, P≤0.001=***, P≤0.0001=****.

### Environmental moisture and dietary protein-to-carbohydrate content dictate life history traits in axenic flies

While the increase in protein and decrease in carbohydrates due to gut bacteria were not surprising based on previous studies showing the influence of bacteria on protein nutrition and carbohydrate utilization (Huang and Douglas, 2015; Keebaugh et al., 2018), we were not expecting such a significant shift in moisture content. We hypothesized that, in addition to altering protein and carbohydrates, bacterial manipulation of moisture in fly diets would have resulting effects on fly life history.

To increase moisture in fly food without diluting the diet we reared axenic flies in conditions of low (27%) or high (85%) relative humidity (RH). Fly vials placed in high humidity were also treated with 300 μL of 1% agar as in Ja *et al.* to further promote water availability (Ja et al., 2009). We found that high humidity delayed development and extended longevity across diets (**Figure 3A-B**). These results are consistent with previous studies showing that increased dietary water content slowed larval development time and lengthened lifespan of flies fed a concentrated, nutrient-rich diet (Hodge, 2001; Ja et al., 2009). Our results showing that this is influenced by the microbiome suggest a novel mechanism by which microbial association can impact *D. melanogaster* physiology.

**Figure 3.**
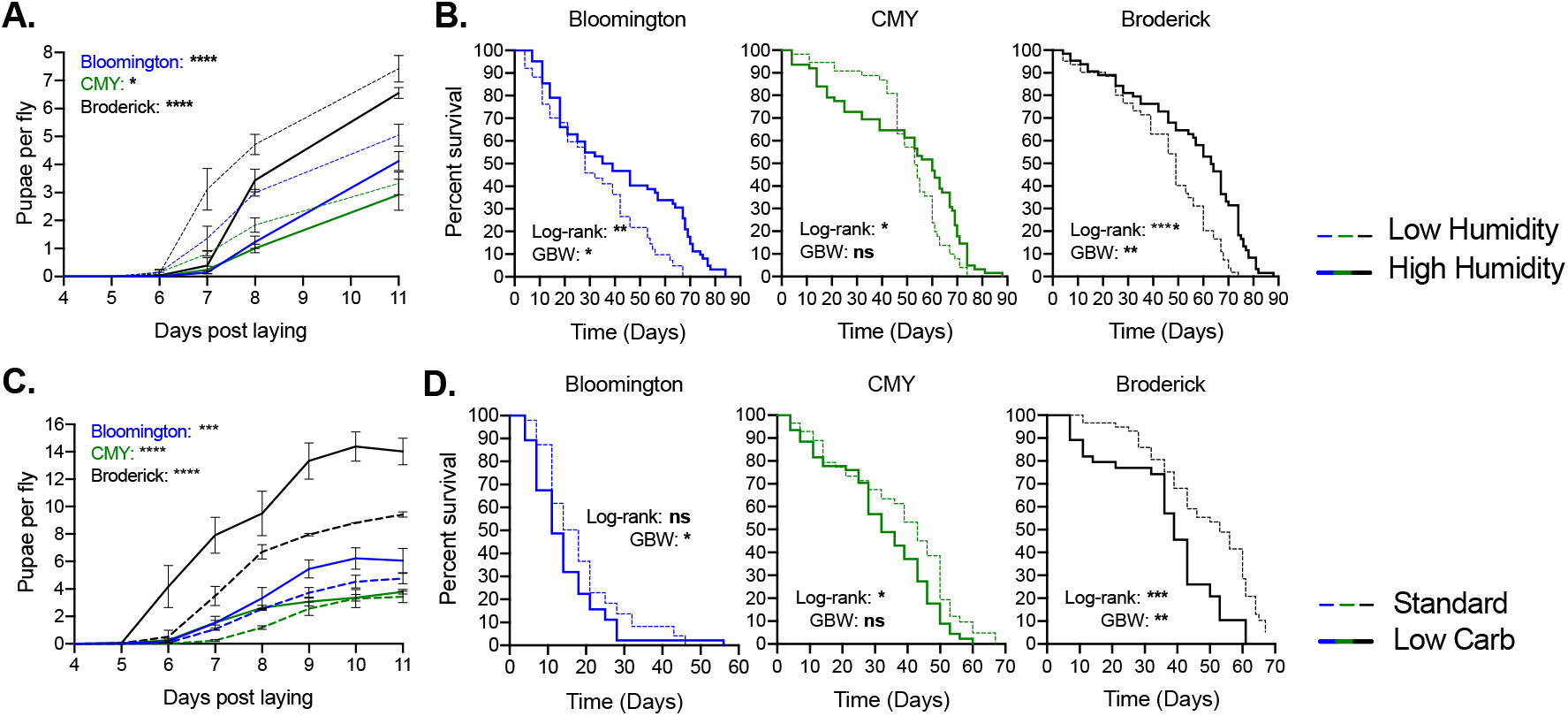
Environmental moisture and dietary protein-to-carbohydrate content dictate life history traits in axenic flies. **A)** Fecundity and **B)** longevity of axenic flies reared on the Bloomington, CMY, and Broderick diets at low (27%) or high (85%) relative humidity (RH). **C)** Fecundity and **D)** longevity of axenic flies reared on standard Bloomington, CMY, and Broderick diets (Standard) or with carbohydrates reduced by 29% (Bloomington), 42% (CMY), or 31% (Broderick) in an attempt to match protein-to-carbohydrate ratio of the top half of each diet with bacteria as determined from **Figure 2** nutritional analyses (Low Carb treatment). Each graph **(A-D)** represents the average of three replicate experiments starting with 18-25 4-day old adult females. Fecundity **(A,C)** is expressed as number of pupae recorded for 11 days divided by number of laying adult females in vial; lines represent mean of replicates ± SEM; statistical analyses were performed by two-way ANOVA with Bonferroni multiple comparisons for Low and High treatments **(A)** or for Standard versus Low Carb treatments **(C)** within each diet. Longevity **(B,D)** is expressed as Kaplan-Meier survival with statistical difference between Low and High humidity **(B)** or between Standard and Low Carb treatments **(D)** on each diet determined via Log-rank and Gehan-Breslow-Wilcoxon (GBW) analyses for the 3 replicate experiments combined. Survival curves for individual replicates are shown in **Figure S2**. Significance is expressed as: P>0.05=ns, P≤0.05=*, P≤0.01=**, P≤0.001=***, P≤0.0001=****.

The observed changes in protein and carbohydrate levels in food with bacteria are reminiscent of dietary restriction studies, in which alteration of protein-to-carbohydrate ratio (P:C) leads to significant effects on life history (Frieling and Roeder, 2020; Partridge et al., 2005). To measure the impact of bacteria-mediated dietary carbohydrate and protein changes, we prepared diets with roughly 30% lower carbohydrate content by decreasing the amount of corn syrup in the Bloomington diet, molasses in the CMY diet, and sucrose in the Broderick diet. Consequently, the resulting “low carbohydrate” diets were composed of similar ratios of protein to carbohydrates (P:C) as the top half of each diet containing bacterial treatment from **Figure 2**(**Table S1**). As shown in dietary restriction studies in which decreased P:C results in reduced fecundity, but extended longevity, increasing P:C through carbohydrate reduction increased fecundity while reducing longevity (**Figure 3C-D**). Feeding rate was measured in young flies on each diet and was found to be unaffected by low carbohydrate conditions on the CMY and Broderick diets, but was slightly increased in Bloomington Low Carb flies compared to control (**Figure S2**). However, feeding rate was noticeably lower in general on the two Bloomington diets compared to the other diets, which may explain the significant amounts of early death on the Bloomington diets (**Figure 3D, Figure S2, Figure S3**).

### Gut bacteria increase dietary tryptophan with resulting impacts on longevity

To further understand bacteria-mediated impacts on protein content in the context of amino acid provisioning, we analyzed the Bloomington diet for amino acid content with and without 14 days of bacterial growth. The only amino acid significantly impacted by bacteria was tryptophan, which increased in the top half of food with bacterial inoculation (**Figure 4A**, see **Figure S4** for complete amino acid profile). We then examined whether increases in tryptophan due to bacteria in food influence host life history traits. We prepared the Bloomington diet as normal (Control, containing 0.24 g/L tryptophan), or supplemented food with tryptophan that either matched bacteria-induced levels (Low Trp, 0.31 g/L total tryptophan) or greatly increased the tryptophan concentration (High Trp, 3.7 g/L total tryptophan), and monitored life history of axenic flies on each treatment. We observed no impact of altered tryptophan on fly fecundity (**Figure 4B**), but for longevity, found that low and high tryptophan, while neither was statistically different from the control, were significantly different from each other, with lifespan reduced on high tryptophan compared to low (**Figure 4C**). The implications of these results are challenging to interpret, as Control survival varied considerably across replicate experiments (**Figure S3**). It is likely that the small difference in tryptophan levels between control and the Low Trp treatment (0.24 g/L compared to 0.31 g/L) is not sufficient enough to visualize a change in lifespan, but it is unclear if the same can be said between Control and High Trp treatments (0.24 g/L compared to 3.7 g/L). Regardless, a significant reduction in lifespan, particularly in flies surviving longer than 30 days, was observed in High Trp compared to Low, suggesting a role for tryptophan in aging on the Bloomington diet.

**Figure 4.**
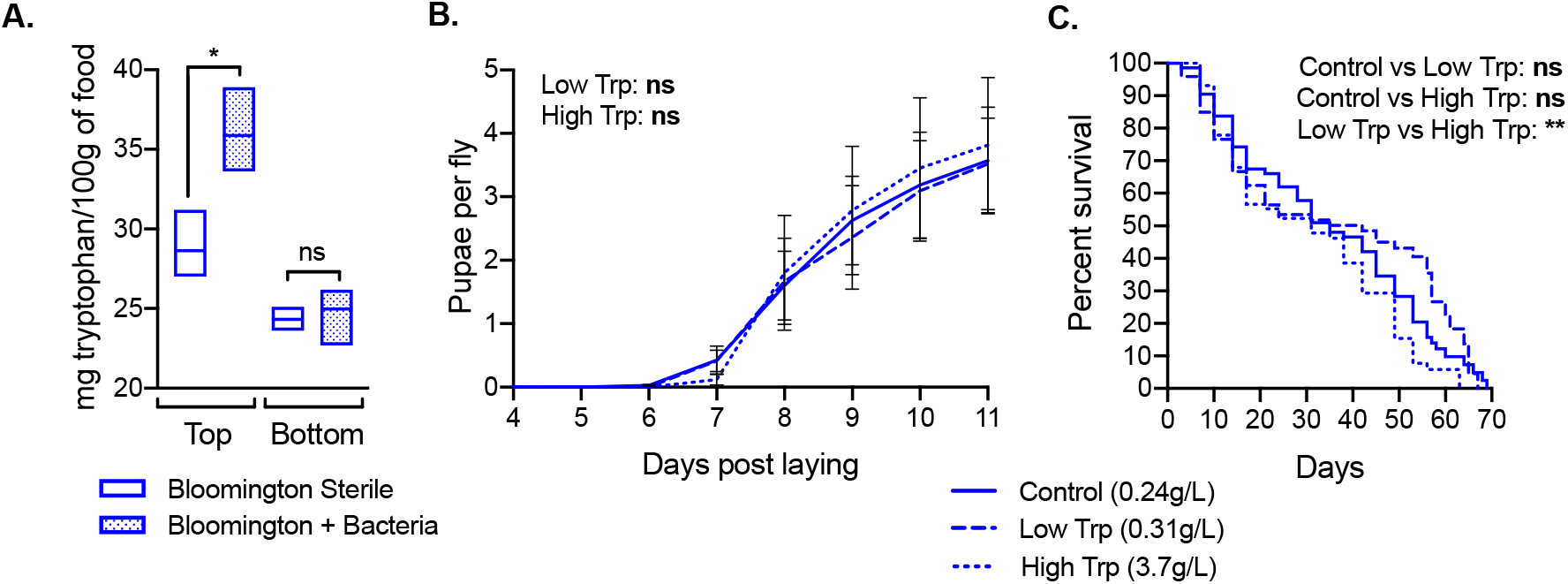
Gut bacteria increase dietary tryptophan with resulting impacts on longevity. **A)** Amount of tryptophan in Bloomington diet 14 days after inoculation with PBS (Sterile) or 10^4^ cells of a 4-species bacterial cocktail (+Bacteria). As in **Figure 2**, the top (~7 mm) and bottom (~7 mm) half of food were analyzed separately. Bars represent minimum and maximum values and mean of 3 biological replicates. Statistical significance of Sterile vs +Bacteria treatments were determined using unpaired two-tailed t-tests. **B)** Fecundity and **C)** longevity of axenic female flies reared on the Bloomington diet prepared as standard (Control, 0.24 g/L total tryptophan), supplemented with 71.5 mg L-tryptophan (Low Trp, 0.31 g/L total tryptophan), or supplemented with 3.5 g L-tryptophan (High Trp, 3.7 g/L total tryptophan). Results are compiled from 3 replicate experiments starting with 25 3-day old females each. Fecundity **(B)** is expressed as number of pupae recorded for 11 days divided by number of laying adult females in vial; lines represent mean ± SEM; statistical analyses performed by two-way ANOVA with Dunnett’s multiple comparisons analysis comparing low and high tryptophan samples to control. Longevity **(C)** is expressed as Kaplan-Meier survival with statistical difference between each treatment determined via Log-rank analysis (Gehan-Breslow-Wilcoxon statistical analysis was not significant). Survival curves for individual replicates are shown in **Figure S2**. Significance is expressed as: P>0.05=ns, P≤0.05=*, P≤0.01=**, P≤0.001=***, P≤0.0001=****.

### Gut bacteria influence host life history differentially based on fly diet

To contextualize our results showing life history impacts of P:C, moisture, and tryptophan, we next addressed how microbes themselves affect physiology on the three standard diets. We found that diet was a crucial factor determining microbial impacts on development time, number of pupae per fly, and fly lifespan. While +Bacteria treatments profoundly increased pupae numbers on the Bloomington and CMY diets, it had no effect on total pupae counts on the Broderick diet (possibly due to a decrease in feeding rate with bacterial treatment; **Figure S2**), however development time to pupation was faster on all three diets with bacterial treatment (**Figure 5A**, most evident at Day 6 in Broderick diet; Day 7 in Bloomington and CMY diets). Lifespan of flies exposed to bacteria was most impacted on the Broderick diet, with +Bacteria conditions reducing median lifespan by about 10 days compared to Sterile controls (**Figure 5B**). While Bloomington flies treated with bacteria also experienced slightly shorter lifespans compared to Sterile, Sterile and +Bacteria CMY flies did not have discernably different survival (**Figure 5B**). No difference in bacterial load or composition (by genus) were observed in flies throughout their lifespan across the three diets (**Figure 5C**), suggesting that diet-specific bacterial effects on fecundity and longevity were not due to differences in bacterial growth, but potentially to microbial utilization and/or provisioning of nutrients or other metabolites that differ based on specific dietary composition.

**Figure 5.**
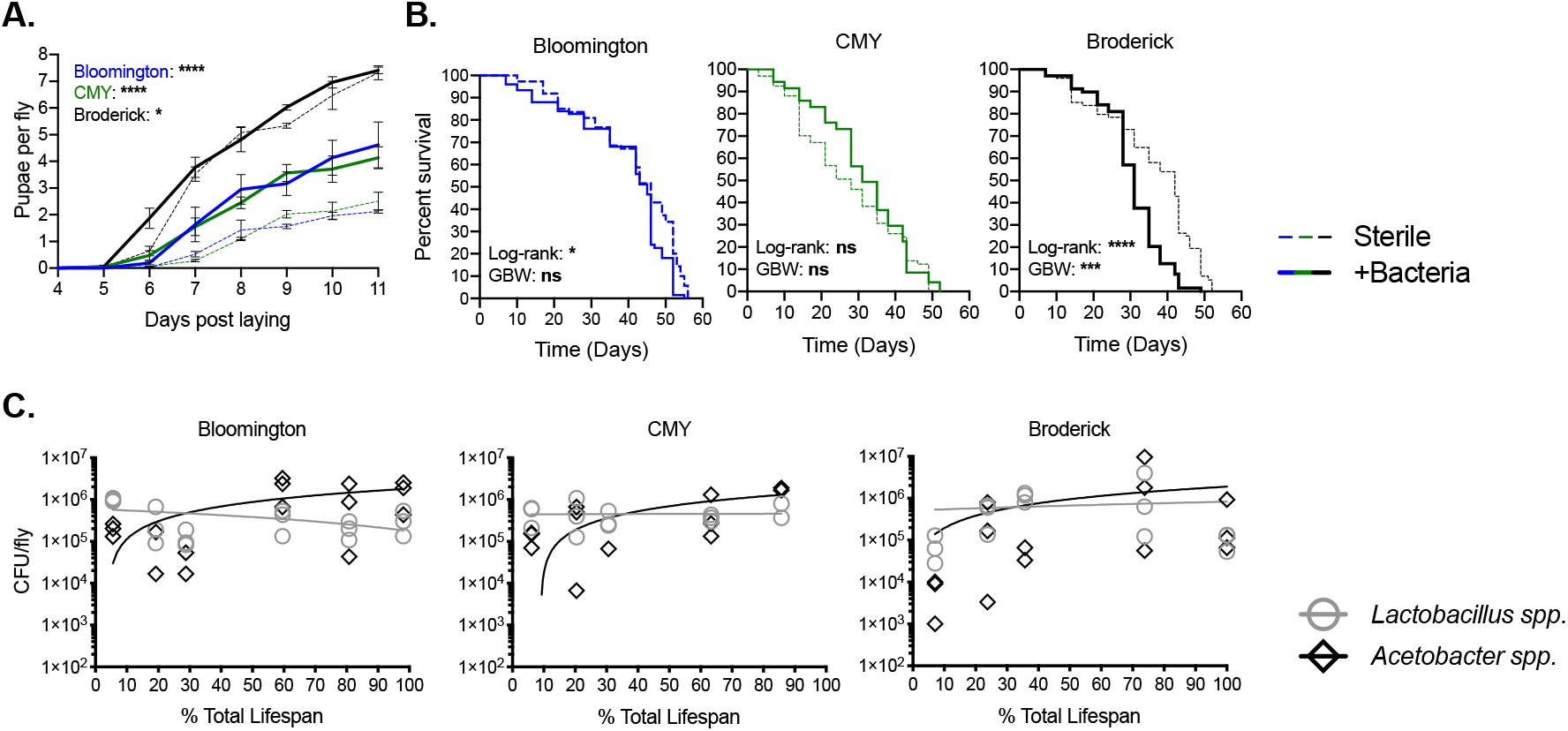
Gut bacteria differentially influence host life history based on fly diet. **A)** Fecundity, **B)** longevity, and **C)** bacterial load of (initially) axenic female flies maintained on the Bloomington, CMY, and Broderick diets supplemented with PBS (Sterile) or 10^4^ bacterial cells (+Bacteria) with each passage to fresh food. Fecundity, longevity, and bacterial load data was obtained from 3 **(A,B)** or 1 **(C)** replicate experiments consisting of 21-25 4-day old adult flies at the experiment start. Fecundity **(A)** is expressed as number of pupae recorded for 11 days divided by number of laying adult females in vial; lines represent mean of replicates ± SEM; statistical analyses were performed by two-way ANOVA with Bonferroni multiple comparisons for Sterile and +Bacteria treatments within each diet. Longevity **(B)** is expressed as Kaplan-Meier survival with statistical difference between Sterile and +Bacteria on each diet determined via Log-rank and Gehan-Breslow-Wilcoxon (GBW) analyses for the 3 replicate experiments combined. Survival curves for individual replicates are shown in **Figure S3**. Bacterial load **(C)** is shown as a combination of *L. plantarum* and *L. brevis* (*Lactobacillus spp*.) or of *A. pasteurianus* and *A. tropicalis* (*Acetobacter spp.)* for individual flies (each point = 1 fly); best-fit lines for each genus were determined via linear regression (no significant difference in *Lactobacillus spp.* or *Acetobacter spp.* when each diet is compared with one another). Significance is expressed as: P>0.05=ns, P≤0.05=*, P≤0.01=**, P≤0.001=***, P≤0.0001=****.

### Gut bacteria impact nutrition in a natural food substrate

Having investigated bacteria-mediated changes in nutrition on three standard laboratory diets, we asked how the observed nutritional changes translated to a natural fly dietary substrate. We analyzed nutritional content of grapes as fresh samples (with PBS) or after inoculation with the 4-species bacterial cocktail after 14 days of incubation. As observed in standard diets (**Figure 1)**, bacterial growth on grapes resulted in a decrease in carbohydrates and increase in moisture; a trend towards higher protein was also observed, but did not meet significance at P=0.05 level (P=0.06, **Figure 6A-C**). Surprisingly, we detected only *Lactobacillus spp.* after incubation, suggesting that the observed nutritional effects, at least on grapes, were mediated primarily by *Lactobacillus*, and were not dependent on co-culture with or contributions from *Acetobacter* (**Figure 6D**). Together, these results confirm that the nutritional relationship between *D. melanogaster* gut microbes and “standard” laboratory fly food is not only reproducible in a natural food source, but also physiologically relevant.

**Figure 6.**
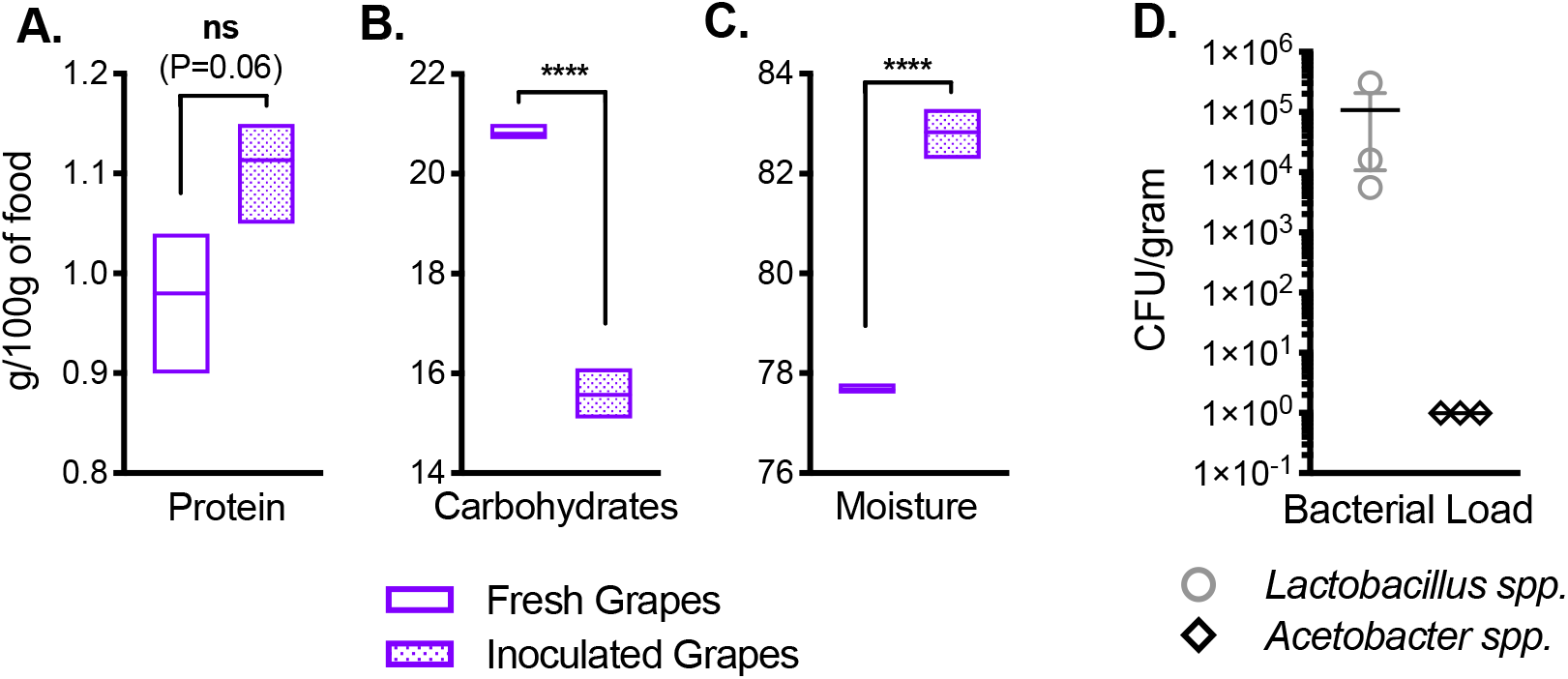
Gut bacteria impact nutrition in a natural food substrate. **A)** Protein, **B)** carbohydrates, **C)** moisture, and **D)** bacterial load in crushed grapes either immediately after inoculation with PBS (Fresh) or 14 days after inoculation with 10^4^ cells of a 4-species bacterial cocktail (Inoculated). Bars in **A-C** represent minimum and maximum values and mean of 3 biological replicates; statistical differences between Fresh and Inoculated treatments were determined using unpaired two-tailed t-tests. Bacterial load **(D)** is expressed as colony forming units (CFU) per gram of food with each point representing an individual replicate; lines and error bars represent mean ± SEM; bacterial growth was not detected on fresh grapes and is not shown. Significance is expressed as: P>0.05=ns, P≤0.05=*, P≤0.01=**, P≤0.001=***, P≤0.0001=****.

## Discussion

Gut microbes can rescue the detrimental effects of flies feeding on diets lacking protein (Keebaugh et al., 2018; Shin et al., 2011), yet reduce lifespan in certain nutritionally rich conditions (Keebaugh et al., 2019). However the nutritional impact of microbiota of fruit flies reared on “standard” laboratory diets or natural dietary environments has not been extensively explored. In this study, we investigated the impact of a representative bacterial community on three standard diets and a natural diet of grapes on the nutritional makeup of food and demonstrate their effects on host-microbe physiology. Because of *D. melanogaster*’s close association with its microbe-rich food source, which heavily dictates the host’s internal microbiome (Chandler et al., 2011), studies of fly diet external to the host provide insight into critical biological processes mediated by the microbiome.

### Nutrient stratification

Growth of bacteria on each diet resulted in decreases in carbohydrates and increases in moisture throughout the food, yet protein was only detectably increased in the top half of food. This shows that microbial growth/activity leads to nutrient heterogeneity in the fly substrate (**Figure 2**). The stratification of protein is correlated with higher bacterial counts in the top half of food, suggesting that bacterial biomass itself contributed to the significant increase in dietary protein. The fact that carbohydrate and moisture levels were constant throughout the food, unlike protein, suggests that higher bacterial biomass on the food surface was not accompanied by increased bacterial metabolism (which would result in sugar consumption and presumably moisture production), possibly due to a “maxing out” of metabolic activity throughout food despite the varying rates of cell division (Liu, 1998). Protein stratification due to bacteria is important to consider in the context of nutrition-dependent development and longevity, as larvae and adults may be exposed to different nutritional environments. Larval density likely also impacts this nutrient stratification, such that increased churning of substrate at the food surface may lead to better mixing of food layers than is seen in tubes lacking (or with fewer numbers of) larvae (Sokolowski et al., 1997).

### Microbiome and dietary plenitude

Dietary restriction studies have identified decreased protein availability as a factor that consistently prolongs lifespan and health span across model organisms (Frieling and Roeder, 2020). In flies, this effect is tied to insulin signaling and the TOR pathway, which are stimulated by both diet and the microbiome (Dobson et al., 2016; Koyama et al., 2014; Shin et al., 2011; Storelli et al., 2011). Keebaugh *et al.* recently showed that live and dead bacteria extended lifespan of protein-deficient flies (Keebaugh et al., 2018), but had less of an effect on high nutrient diets (Keebaugh et al., 2019). These studies identify protein availability as a major factor defining the host-microbe relationship, but acknowledge that overall impacts of microbes on lifespan are likely a combination of nutritional, host, and microbial factors. We show that the microbiome itself alters the nutritional composition of diet to generate conditions of dietary plenitude, specifically an excess of protein accompanied by decreased carbohydrates, and that this occurs to a similar extent on three standard diets despite different nutritional compositions. While these dietary changes result in developmental and longevity phenotypes characteristic of both high P:C diets and flies reared with microbes on the Bloomington and Broderick diets, the relationship is not as obvious with regard to longevity on the CMY diet. No significant difference in lifespan was observed between Sterile and +Bacteria treatments on CMY food, suggesting further nutrient-mediated effects of bacteria on life history independent of protein and carbohydrates.

A possible factor influencing the different impacts of bacteria on life history between diets is micronutrients, which differ between diets based on specific nutrient composition (Cantarel et al., 2012; Lesperance and Broderick, 2020b; Li et al., 2011; Muegge et al., 2011; Turnbaugh et al., 2006). While we did not directly test the effect of micronutrient differences between the three standard diets, we can make some predictions based on their unique dietary components. For example, the CMY diet contains molasses, which is rich in choline (13.3 mg per 100 g of molasses; Lesperance and Broderick, 2020b). Choline is a compound known both to be metabolized by gut bacteria (Romano et al., 2015) and impact *D. melanogaster* growth and development (Geer and Vovis, 1965), making it just one of many possible candidate micronutrients that could conceivably modulate microbiome-diet-host relationships.

### Microbiome as a regulator of tryptophan metabolism

In addition to micronutrients, amino acids are likely to impact host biology in different ways based on host-microbe, host-diet, and microbe-diet relationships (Consuegra et al., 2020; Grandison et al., 2009; Judd et al., 2018; Manière et al., 2020; Roager and Licht, 2018; Yamada et al., 2015). Our analysis showed that tryptophan was significantly increased with bacterial growth and, when administered in the diet to axenic flies, impacted fly lifespan. As a precursor for the hormone serotonin, tryptophan’s connection with the microbiome is the subject of many recent investigations of the gut-brain axis-that is, the role of the microbiome in cognitive function and as a factor in disorders such as anxiety and depression (reviewed in O’Mahony et al., 2015 and Gao et al., 2018). Studies in flies and rodents exploring the function of tryptophan in host physiology independent of the microbiome have identified a role for the amino acid in aging-related pathologies, some even drawing links between tryptophan deficiency and dietary restriction (Breda et al., 2016; Burnet and Sang, 1968; De Marte and Enesco, 1986; Ooka et al., 1988; Oxenkrug et al., 2011; Segall and Timiras, 1976). As in humans, tryptophan is an essential amino acid for *D. melanogaster*, meaning the amino acid must be acquired from dietary sources for complete nutrition (Sang and King, 1961). Yet, tryptophan is relatively scarce in nature and metabolically expensive for plants and microbes to produce (Radwanski and Last, 1995), highlighting microbiome-mediated tryptophan metabolism as an important aspect of *D. melanogaster* host-microbe physiology. It is unclear if any of the gut microbes utilized in this study biosynthesize tryptophan or if it is extracted from dietary proteins, but not utilized (Consuegra et al., 2020; Macfarlane et al., 1986; Mcfarlane and Allison, 1986), so additional work exploring microbial regulation of tryptophan in *D. melanogaster* will be of great interest.

### Bacteria-derived moisture

Our study is the first to link growth of gut bacteria on the fly diet to significant increases in dietary moisture, but it is not yet clear how such changes contribute to microbe-mediated host physiology. Our results suggest that the relationship between bacteria, dietary moisture, and life history is complex, as we observed the microbiome as a factor reducing development time and shortening lifespan on two of three diets, contrary to what we observed with increased environmental moisture. It is likely that factors modulating the impact of the microbiome on life history are multi-faceted, and microbial contributions to dietary moisture may not be significant enough *in vivo* to greatly affect physiology.

With regard to behavioral physiology, increased moisture likely contributes to the attractiveness of fermented foods to the fly. “*Drosophila,”* derived from Greek, means “dew-loving,” indicating a long-standing appreciation for the necessity of moist environments for fly habitation (Perveen, 2018). Sayeed *et al.* reported hygrosensory behavior in *D. melanogaster*, confirming that moisture is specifically sought out by the fly (Sayeed and Benzer, 1996). Previous work investigating attractive compounds produced by gut microbes has focused primarily on metabolites and odorants (Ai et al., 2010; Fischer et al., 2017; Joseph et al., 2009; Seong et al., 2020), but the microbiome has been underappreciated as a source of attractive moisture content in the dietary substrate. It’s also not clear how the microbiome increases food moisture content, but it’s possible this could be directly as a byproduct of metabolism or indirectly through reduced evaporation from the food surface, for example via biofilms or surfactants (Fechtner et al., 2011; Lennon and Lehmkuhl, 2016).

### Physiologic relevance

We conclude our study by confirming that the protein, carbohydrate, and moisture changes in standard fly diets translate to a natural fly dietary substrate inoculated with bacteria. Because we also confirmed that inoculating standard diets via gnotobiotic flies results in similar nutritional changes as culture inoculation, we expect that the observed nutritional changes occur in fly-associated food substrates in nature. It has long been appreciated that microbes including *Lactobacillus*, *Acetobacter*, and yeasts are crucial for promoting fly association with natural dietary sources including fruit (reviewed extensively in Broderick and Lemaitre, 2012). Our study suggests, that in modifying substrate to provide a plentiful, moist, nutritional environment, *D. melanogaster*-associated microbes create an attractive, hospitable niche supportive of larval development and adult homeostasis. Future work exploring whether micronutrient differences in diet or specific microbiome compositions vary microbe-mediated effects on the host will greatly expand our understanding of fly nutritional physiology and ecology.

## Acknowledgements

We thank Alexander Barron, Madison Condon, and Rose Dziedzic for critical editing of the manuscript and moral support. Funding was provided by the National Institutes of Health [R35GM128871] and the University of Connecticut. Stocks obtained from the Bloomington *Drosophila* Stock Center (NIH P40OD018537) were used in this study. Figures were created in GraphPad Prism and BioRender.com.

## Author Contributions

Conceptualization, D.N.A.L. and N.A.B.; Methodology, D.N.A.L. and N.A.B.; Formal Analysis, D.N.A.L.; Investigation, D.N.A.L.; Writing – Original Draft, D.N.A.L.; Writing – Review & Editing, D.N.A.L. and N.A.B.; Visualization, D.N.A.L.; Supervision, N.A.B.; Funding Acquisition, N.A.B.

## Methods

### Fly diets

Diets used for nutritional analyses included the Bloomington Standard Nutri-Fly formulation (Genesee Scientific, 66-113), Bloomington Molasses Nutri-Fly formulation (Genesee Scientific, 66-116), and the Broderick Standard diet (**Table 1**). For life history experiments with altered carbohydrates, Bloomington Standard, Bloomington Cornmeal-Molasses-Yeast, and Broderick Standard diets were made from scratch with reduced amounts of corn syrup, molasses, or sucrose, respectively (**Table 1**). Life history experiments with diets containing altered tryptophan content were done using the Bloomington Standard Nutri-Fly formulation with 71.5 mg (low) or 3.5 g (high) of L-tryptophan (Sigma T8941) added as per Breda et al., 2016. Fly food was prepared by boiling 1L of distilled water, stirring in dry ingredients, and allowing food to cook for 20 minutes. Methyl paraben was added last before mixing food via immersion blender. Diets were transferred to wide vials (10mL) or 6oz bottles (50mL). Vials/bottles were plugged and autoclaved for 20 minutes at 121°C.

**Table 1.** Description of experimental diets.

| Diet | Components per 1L Water & Source |
| --- | --- |
| Bloomington Standard (Nutri-Fly) | Genesee Scientific 66-113<br>Nutri-Fly BF<br>Yellow cornmeal, agar (Type II), corn syrup solids, inactive nutritional yeast, soy flour<br>4.4 mL propionic acid added during cooking |
| Bloomington Molasses (CMY) (Nutri-Fly) | Genesee Scientific 66-116<br>Nutri-Fly MF<br>Cornmeal, agar (Type II), molasses solids, inactive nutritional yeast<br>1.4 g methyl paraben in 10mL 100% ethanol added during cooking |
| Broderick Standard | 50 g inactive dry yeast<br>70 g yellow cornmeal<br>6 g agar<br>40 g sucrose<br>1.25 g methyl paraben in 5mL 100% ethanol |
| Bloomington<br>Standard<br>(Homemade) | 15.9 g inactive dry yeast<br>67 g yellow cornmeal<br>9.2 g soy flour<br>5.29 g agar<br>102 mL corn syrup<br>4.4 mL propionic acid<br>https://bdsc.indiana.edu/information/recipes/bloomfood.html<br>(Recipe makes 42.5 L, so all ingredient amounts were divided by 42.5 to make 1 L) |
| Bloomington<br>CMY<br>(Homemade) | 12.4 g inactive dry yeast<br>61.3 g yellow cornmeal<br>6.01 g agar<br>75.2 mL molasses<br>1.4 g methyl paraben in 10mL 100% ethanol<br>https://bdsc.indiana.edu/information/recipes/molassesfood.html<br>(For this recipe, values were divided by 2.66 to make 1 L) |
| Bloomington<br>Low<br>Carbohydrate | 15.9 g inactive dry yeast<br>67 g yellow cornmeal<br>9.2 g soy flour<br>5.29 g agar<br>41 mL corn syrup<br>4.4 mL propionic acid |
| CMY Low<br>Carbohydrate | 12.4 g inactive dry yeast<br>61.3 g yellow cornmeal<br>6.01 g agar<br>15 mL molasses<br>1.4 g methyl paraben in 10mL 100% ethanol |
| Broderick<br>Low<br>Carbohydrate | 50 g inactive dry yeast<br>70 g yellow cornmeal<br>6 g agar<br>0 g sucrose<br>1.25 g methyl paraben in 5mL 100% ethanol |

### Preparation of inoculum

*Lactobacillus plantarum, Lactobacillus brevis,* and *Acetobacter pasteurianus* strains used were previously isolated from lab-reared *Drosophila melanogaster* by the Broderick lab. *Acetobacter tropicalis* was obtained from John Chaston (Judd et al., 2018). Liquid cultures were prepared as follows shaking at 200 rpm: *Acetobacter* strains: grown 1 day in a 1:1 mixture of Man, Rogosa, and Sharpe (MRS) and mannitol broth, *Lactobacillus* strains: grown for 1 day in MRS broth. Cultures were centrifuged for 20 minutes at 4°C at 3428 x g, supernatant was removed, and cells were resuspended in sterile PBS. Each bacterium was then quantified using a hemocytometer and set to a concentration of 10^4^ cells per 500 μL in PBS. Each strain was then combined at 1:1:1:1 and 500 μL of the bacterial mix were pipetted onto fly food for nutritional analysis experiments; sterile controls were treated with 500 μL of PBS.

### Axenic and gnotobiotic flies

All flies used in this study were from the Oregon-R background (which contains *Wolbachia*) from the Bloomington *Drosophila* Stock Center (Bloomington, IN). Flies were maintained in a 25°C 12h:12h light:dark incubator in ambient humidity (around 27% RH). Axenic flies were generated by letting flies lay eggs overnight on grape juice agar plates smeared with yeast paste, scraping eggs into cell strainer baskets using swabs, and applying 10% bleach. Once the chorion was visually confirmed to have disappeared, eggs were washed with sterile water and ethanol before being transferred via pipette to sterile fly food for development. Bacterial inoculum was prepared as described above and 150 μL of the 4-species cocktail was fed to axenic adults for 4 days (one initial inoculum) before 20 male and 25 female flies were used to inoculate experimental diets for nutritional tests. At the time of inoculation via gnotobiotic fly, subsets of flies were homogenized and plated on MRS agar to determine *in vivo* bacterial counts.

### Nutritional analyses

Experimental diets in 6 oz fly bottles were treated as follows and incubated at 25°C for the specified time period. Culture inoculation for whole sample analysis: inoculated directly with PBS or bacteria and incubated 14 days **(Figure S6A, Figure 1A-D)**. Gnotobiotic inoculation for whole sample analysis: subjected to axenic (control) or gnotobiotic flies for 4 days and incubated without flies an additional 10 days **(Figure S6B, Figure 1E-H)**. Culture and embryo inoculation for top versus bottom analysis: treated with 32 μL sterile embryos (as per Linford et al., 2013) and direct inoculation of PBS/bacteria with 11 days of incubation **(Figure S6C, Figure 2)**. Culture inoculation for top versus bottom amino acid analysis: inoculated directly with PBS or bacteria and incubated 14 days **(Figure S6D, Figure 4)**. For top versus bottom experiments, the food line was measured after the incubation period, the bottle was cut open using a sterile razor, and the top and bottom halves of the food were collected separately using autoclaved weigh boats. Fresh red grapes were washed with water and crushed with a sterile gloved hand and 50 grams were distributed into sterile beakers. Grapes were inoculated with 500 μL of PBS (Fresh) or the 4-species cocktail (Inoculated) as in **Figure S6A**, covered with sterile aluminum foil, and collected either immediately (Fresh) or after 14 days of incubation at 25°C (Inoculated).

Food samples were collected into 50 mL conicals and analyzed by Eurofins Food Integrity & Innovations (Madison, WI) for protein, carbohydrates, fat, ash, moisture, caloric content, and in one experiment, amino acids.

At the time of food collection post-incubation, 0.5 g of food (mixed well in conical) was collected and homogenized in PBS, serially diluted, and plated on MRS agar to quantify *Lactobacillus* and *Acetobacter* by genus.

### Fecundity and lifespan experiments

Axenic flies were generated as described above and adapted to the diets tested for at least two weeks prior to beginning longevity and fecundity analysis. 3-4 day old adult females were collected (18-25 flies per vial). Vials were flipped twice per week for the duration of fly lifespan. Fecundity was recorded as the number of pupae that developed each day for the first 11 days following the experiment start. Longevity was recoded as number of dead flies twice per week at the experiment start and more frequently toward the end of fly lifespan. Vials were kept at 25°C with ambient (~27%) relative humidity except for the high humidity test group, which was kept in a 25°C 12h:12h light:dark incubator set to 85% RH. Vials placed in high humidity were also supplemented with 300 μL of 1% agar on the side of the tube as in Ja et al., 2009.

### Feeding Score

Axenic flies that were adapted to experimental diets were placed on appropriate diets (without starving) in which 0.1% erioglaucine disodium salt (final concentration in food) had been thoroughly incorporated and treated as previously described for each test (i.e. inoculated with PBS, bacteria, or nothing). 10-20 female flies (3-4 days old) fed for two hours at 25°C at the same time of day for each replicate experiment and guts from 5 randomly chosen flies per replicate per treatment were dissected. Qualitative feeding scores were assigned to dissected guts by multiplying the intensity of blue coloring (determined visually) by the percentage of the gut exhibiting blue color.

### Data analysis and statistical tests

T-tests comparing Sterile and +Bacteria treated food and feeding score were performed using R. Two-way ANOVAs with Bonferroni multiple comparisons were performed for bacterial load analyses and pupae counts using GraphPad Prism. Differences in survival between treatments on each diet were analyzed using Kaplan-Meier survival curves and Log-rank and Gehan-Breslow-Wilcoxon (GBW) statistical tests in GraphPad Prism; both statistical tests were used to show overall survival differences (Log-rank) and differences driven by early death, as GBW analysis weights early time point more heavily than late. Linear regression comparing bacterial load over the course of fly lifespan was performed in GraphPad Prism. Significance is expressed as: P>0.05=ns, P≤0.05=*, P≤0.01=**, P≤0.001=***, P≤0.0001=****.

