## Supplemental Information for "Gut bacteria mediate nutrient availability *in Drosophila* diets"

Figure S1.

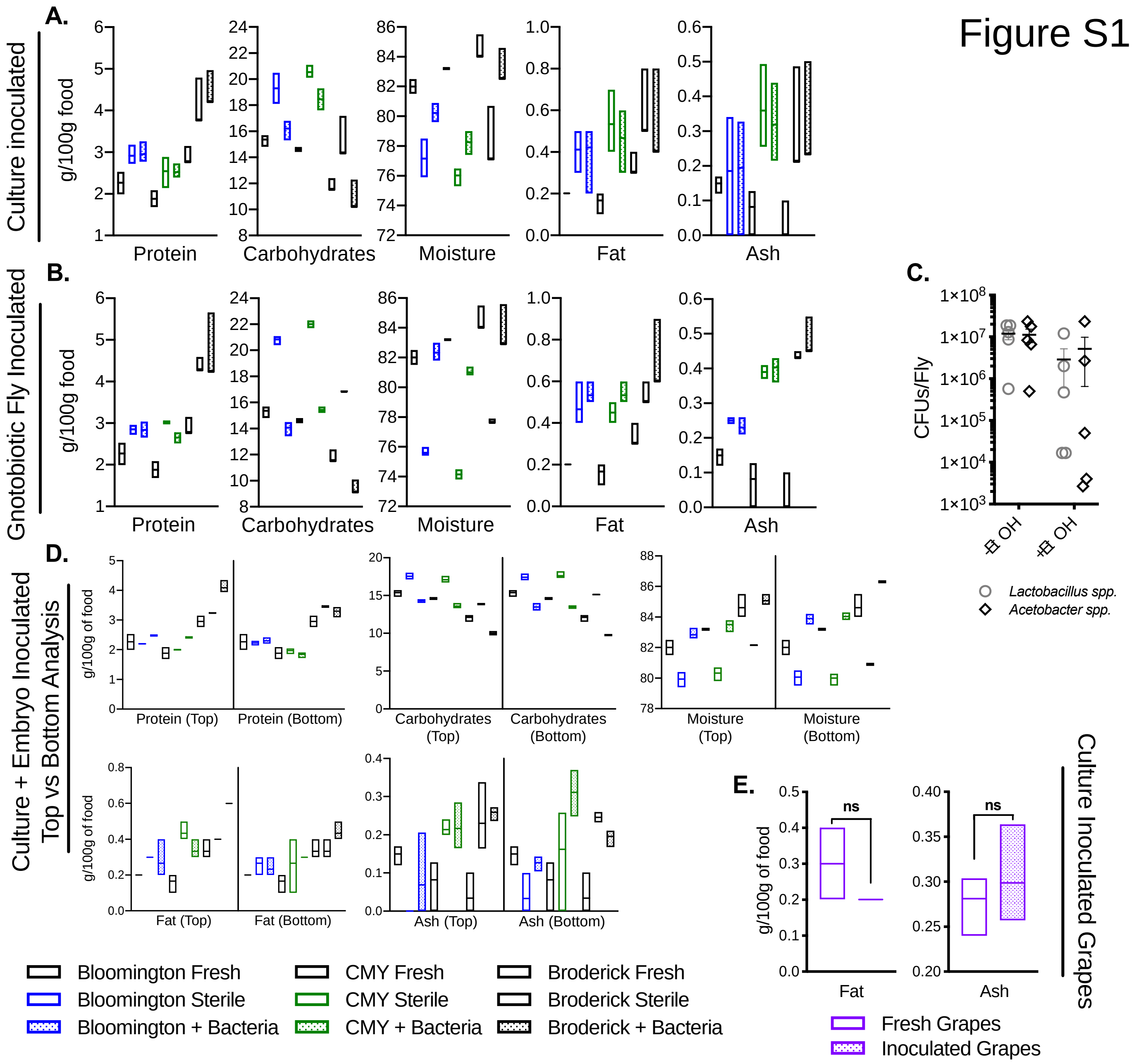


F.

| Experiment | Diet | Nutritional Test | | | | |
|---|---|---|---|---|---|---|
| | | Protein | Carbs | Moisture | Fat | Ash |
| Culture Inoculated | Bloomington | ns | **** | **** | ns | ns |
| | CMY | ns | *** | *** | ns | ns |
| | Broderick | ns | **** | ** | ns | ns |
| Gnotobiotic Fly Inoculated | Bloomington | ns | **** | **** | ns | ns |
| | CMY | * | *** | *** | ns | ns |
| | Broderick | ns | **** | ** | ns | ns |
| Culture + Embryo Inoculated | Bloomington (Top) | *** | *** | ** | ns | ns |
| | CMY (Top) | *** | *** | *** | ns | ns |
| | Broderick (Top) | ** | **** | *** | ns | ns |
| | Bloomington (Bottom) | ns | *** | *** | ns | ns |
| | CMY (Bottom) | ns | *** | *** | ns | ns |
| | Broderick (Bottom) | ns | **** | **** | ns | * |
| | Bloomington (Sterile) | ns | ns | ns | ns | ns |
| | CMY (Sterile) | ns | ns | ns | ns | ns |
| | Broderick (Sterile) | ** | *** | **** | ns | ns |
| | Bloomington (+Bacteria) | ns (0.06) | ns (0.06) | * | ns | ns |
| | CMY (+Bacteria) | *** | ns | ns | ns | ns |
| | Broderick (+Bacteria) | ** | ns | ** | ns | * |

**Figure S1. Raw nutritional values for protein, carbohydrates, moisture, fat, and ash with or without bacterial inoculation**. **A)** Raw nutritional values for Bloomington, CMY, and Broderick diets sampled fresh and14 days after direct inoculation with PBS (Sterile) or $10^4$ cells of a 4-species bacterial cocktail (+Bacteria). **B)** Raw nutritional values for Bloomington, CMY, and Broderick diets sampled fresh and 14 days after inoculation via axenic flies (Sterile) or gnotobiotic flies previously fed $10^4$ cells of a 4-species bacterial cocktail (+Bacteria). **C)** Bacterial load (in colony forming units per fly) in gnotobiotic flies used to inoculate food in **(B)**; CFUs were determined both in ethanol-sterilized (+EtOH) flies and flies not sterilized, as would be the case when flies were used to inoculate diets (-EtOH); each point represents an individual replicate; lines and error bars represent mean ± SEM. **D)** Raw nutritional values for Bloomington, CMY, and Broderick diets sampled fresh and 11 days after inoculation with sterile embryos and PBS (Sterile) or $10^4$ cells of a 4-species bacterial cocktail (+Bacteria); top and bottom half of food were analyzed as separate samples. **E)** Fat and Ash content in crushed grapes either immediately after inoculation with PBS (Fresh) or 14 days after inoculation with $10^4$ cells of a 4-species bacterial cocktail (Inoculated). Statistical differences between Fresh and Inoculated treatments determined via unpaired two-tailed t-test. Bars **(A,B,D)** represent minimum and maximum values and mean of 9 **(A)** or 3 **(B,D,E)** biological replicates; statistical differences between Sterile and +Bacteria treatments within each diet no shading) or between Top and Bottom samples within treatment groups (shaded cells) were determined using unpaired two-tailed t-tests (significance for **A,B,D** shown in **(F)**). Significance is expressed as: $P>0.05$=ns, $P\leq0.05$=*, $P\leq0.01$=**, $P\leq0.001$=***, $P\leq0.0001$=****.

Figure S2.

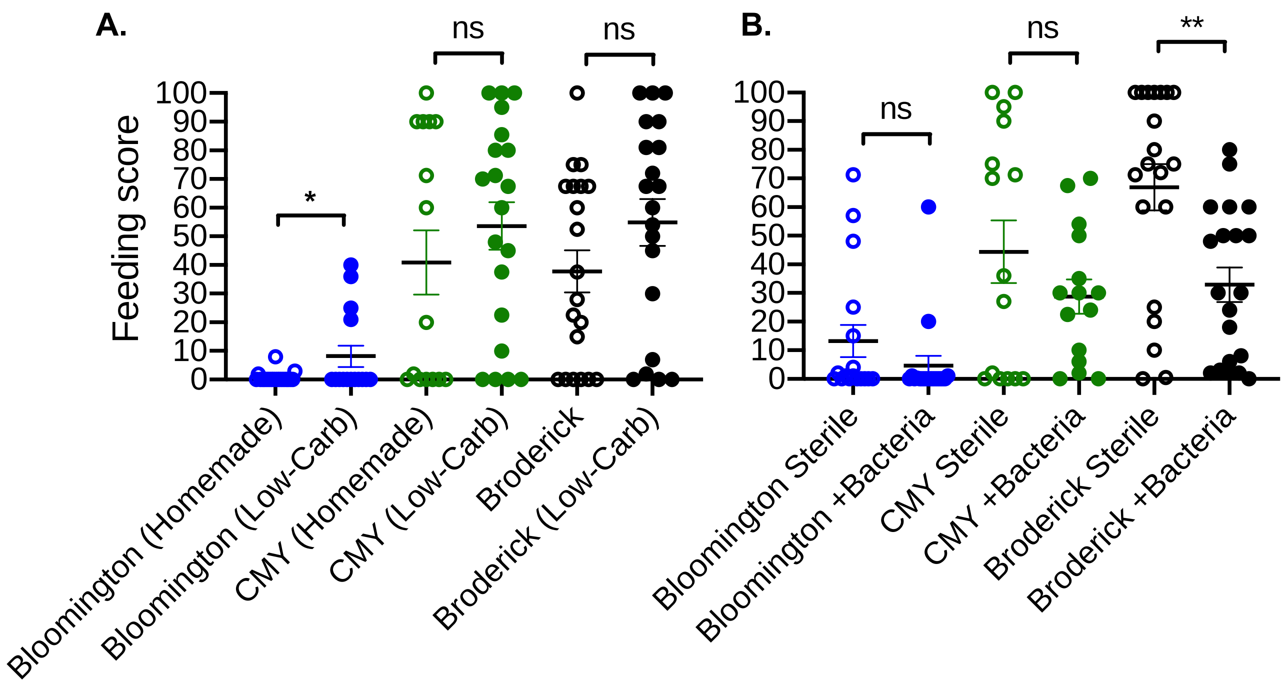


**Figure S2. Feeding rate analysis. A)** Feeding score of flies reared on homemade and low carbohydrate diets. **B)** Feeding score of flies reared on Bloomington (Nutri-fly version), CMY (Nutri-fly version), and Broderick diets supplemented with PBS (Sterile) or the 4-species bacterial cocktail (+Bacteria). Feeding score is a qualitative metric determined by placing 10-20 females (3-4 days old) on food containing 0.1% erioglaucine disodium salt and appropriate treatments, dissecting fly guts (randomly selected from population) after two hours of feeding, and visually scoring the intensity of blue color with in the gut and the percentage of the gut containing coloring. Intensity and length of blue in gut are multiplied to obtain a single feeding score. Data is compiled from 4 replicate experiments randomly sampling 5 flies each; points represent individual flies; lines and error bars represent mean ± SEM. Statistical significance of treatments within each diet determined by unpaired two-tailed t-tests. Significance is expressed as: $P>0.05$=ns, $P\leq0.05$=*, $P\leq0.01$=**, $P\leq0.001$=***, $P\leq0.0001$=****.

### Figure S3.

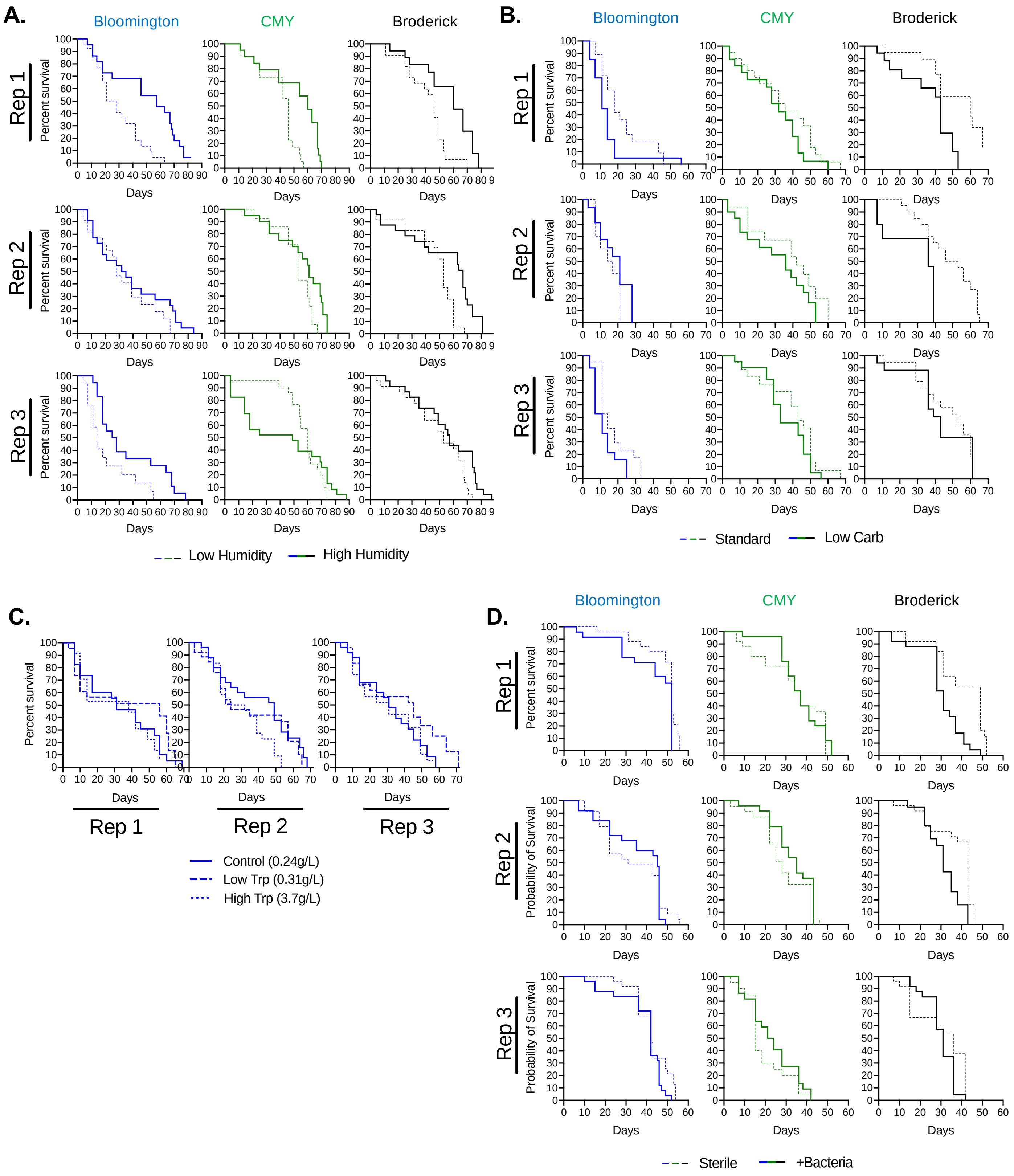

A.
Bloomington
CMY
Broderick
Rep 1
Rep 2
Rep 3
Percent survival
Days
Low Humidity
High Humidity
B.
Bloomington
CMY
Broderick
Rep 1
Rep 2
Rep 3
Percent survival
Days
Standard
Low Carb
C.
Percent survival
Days
Rep 1
Rep 2
Rep 3
Control (0.24g/L)
Low Trp (0.31g/L)
High Trp (3.7g/L)
D.
Bloomington
CMY
Broderick
Rep 1
Rep 2
Rep 3
Percent survival
Probability of Survival
Days
Sterile
+Bacteria

**Figure S3. Individual replicates of survival analyses of flies reared under conditions of altered humidity, dietary carbohydrates, dietary tryptophan, or microbiome. A)** Longevity of axenic female flies maintained on the Bloomington, CMY, and Broderick diets at Low (27%) relative humidity (RH) or High (85% RH) humidity for the duration of the experiment; each replicate longevity experiment began with 18-25 4-day old adult flies. **B)** Longevity of axenic flies reared on standard Bloomington, CMY, and Broderick diets (Standard) or with carbohydrates reduced by 29% (Bloomington), 42% (CMY), or 31% (Broderick) to match protein:carbohydrate ratio of each diet with bacteria as determined from **Figure 2** top half nutritional analyses (Low Carb); each replicate longevity experiment began with 18-25 4-day old adult flies. **C)** Longevity of axenic female flies reared on the Bloomington diet prepared as standard (Control, 0.24 g/L total tryptophan), supplemented with 71.5 mg L-tryptophan (Low Trp, 0.31 g/L total tryptophan), or supplemented with 3.5 g L-tryptophan (High Trp, 3.7 g/L total tryptophan); each replicate longevity experiment began with 25 3-day old adult flies. **D)** Longevity of axenic female flies maintained on the Bloomington, CMY, and Broderick diets supplemented with PBS (Sterile) or $10^4$ bacterial cells (+Bacteria) with each passage to fresh food; each replicate longevity experiment began with 21-25 4-day old adult flies. Data are plotted as Kaplan-Meier survival curves.

### Figure S4.

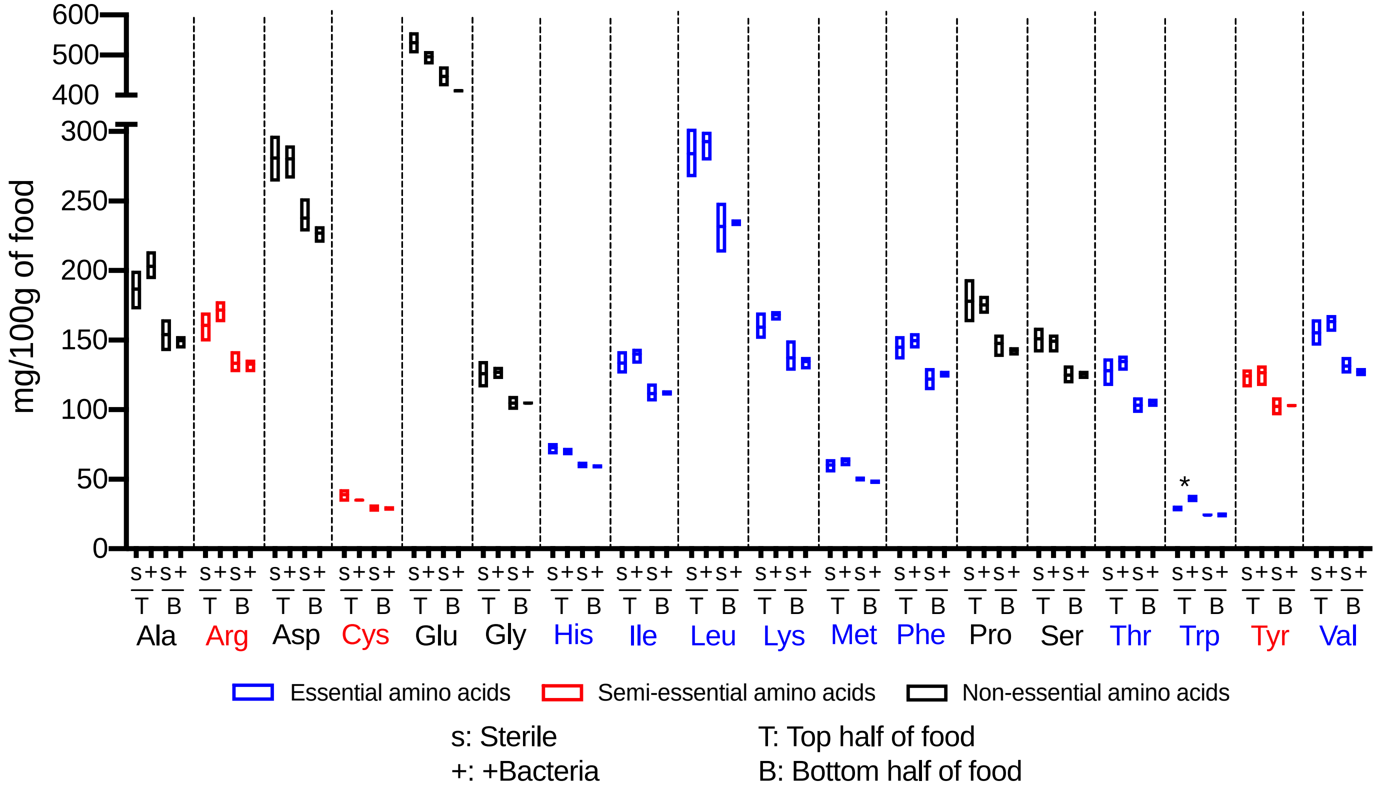


**Figure S4. Tryptophan is the only amino acid detectably impacted by *Drosophila* gut bacteria.** Amino acid content in mg/100g of food in Bloomington diet 14 days after inoculation with PBS (Sterile, s) or 10₄ cells of a 4-species bacterial cocktail (+Bacteria, +); top (T) and bottom (B) half of food were analyzed as separate samples. Bars represent minimum and maximum values and mean of 3 biological replicates; statistical differences between Sterile and +Bacteria treatments within top or bottom for each amino acid were determined using unpaired two-tailed t-tests. Significance is expressed as: P>0.05=ns, P≤0.05=*, P≤0.01=**, P≤0.001=***, P≤0.0001=****.

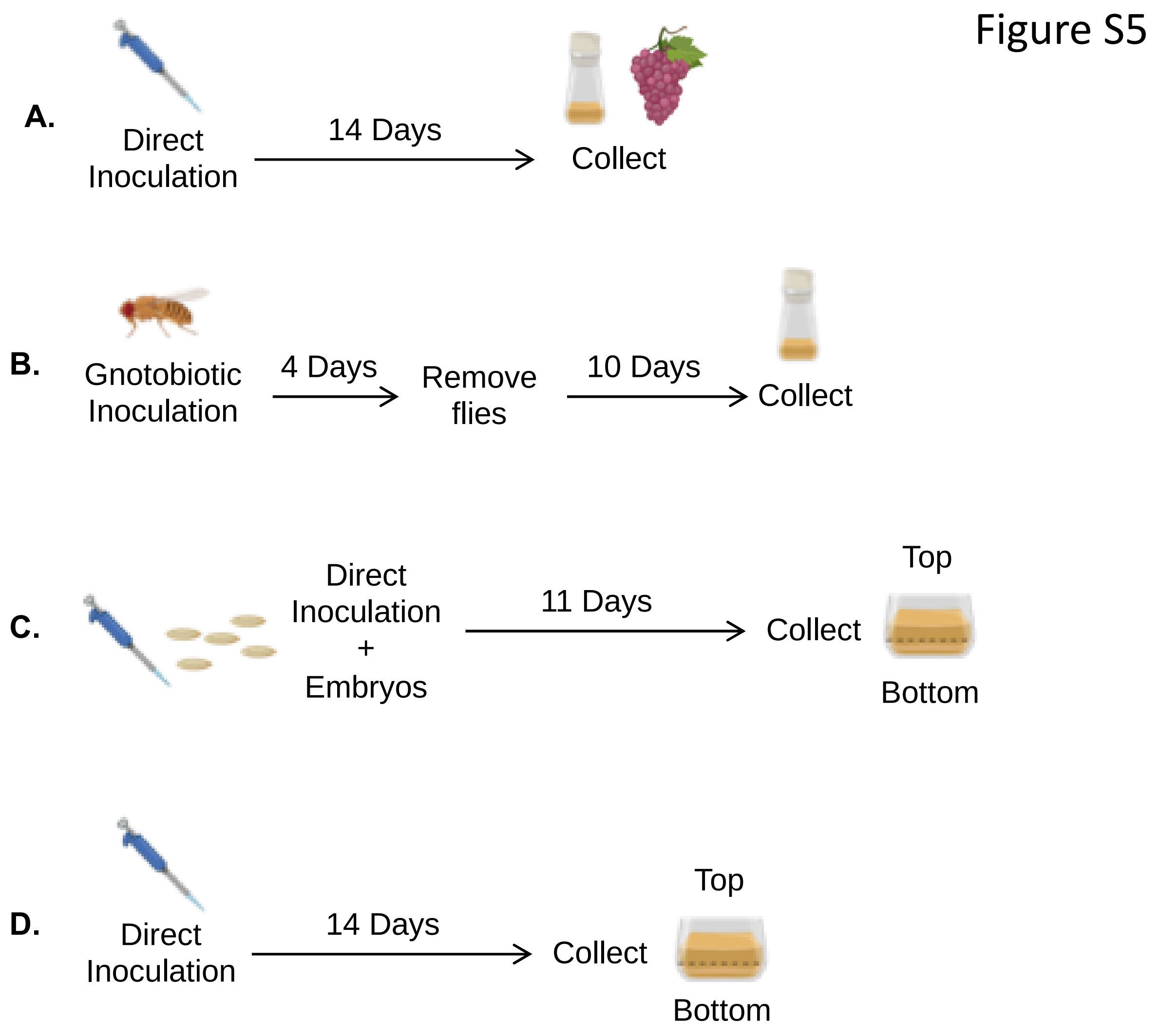


**Figure S5. Experimental design schematics for nutritional content experiments. A)** Culture inoculation method for data shown in **Figure 1A-D**, **Figure S1A**, **Figure 6**, and **Figure S1E**. **B)** Gnotobiotic inoculation method for data shown in **Figure 1E-H** and **Figure S1B-C**. **C)** Inoculation method for data shown in **Figure 2** and **Figure S1D**. **D)** Culture inoculation method for data shown in **Figure 4A** and **Figure S4**.

| Diet | Sterile P:C | +Bacteria P:C |
|---|---|---|
| Bloomington | 0.13 | 0.17 |
| CMY | 0.12 | 0.18 |
| Broderick | 0.23 | 0.41 |

**Table S1. Dietary protein-to-carbohydrate ratios in absence or presence of bacteria.** Comparisons of protein-to-carbohydrate (P:C) ratios in Bloomington, CMY, and Broderick diets 11 days after inoculation with sterile embryos and PBS (Sterile) or $10^4$ cells of a 4-species bacterial cocktail (+Bacteria). Data represents raw protein divided by raw carbohydrate values from top half of food only (see **Figure S1** for raw data).
